# Are Deep Learning Structural Models Sufficiently Accurate for Virtual Screening? Application of Docking Algorithms to AlphaFold2 Predicted Structures

**DOI:** 10.1101/2022.08.18.504412

**Authors:** Anna M. Díaz-Rovira, Helena Martín, Thijs Beuming, Lucía Díaz, Victor Guallar, Soumya S. Ray

## Abstract

Machine learning protein structure prediction, such as RosettaFold and AlphaFold2, have impacted the structural biology field, raising a fair amount of discussion around its potential role in drug discovery. While we find some preliminary studies addressing the usage of these models in virtual screening, none of them focus on the prospect of hit-finding in a real-world virtual screen with a target with low sequence identity. In order to address this, we have developed an AlphaFiold2 version where we exclude all structural templates with more than 30% sequence identity. In a previous study, we used those models in conjunction with state of the art free energy perturbation methods. In this work we focus on using them in rigid receptor ligand docking. Our results indicate that using out-of-the-box Alphafold2 models is not an ideal scenario; one might think in including some post processing modeling to drive the binding site into a more realistic holo target model.

## Introduction

We are witnessing a revolution in scientific methodologies; an avalanche of improved machine learning (ML) techniques is changing scientists’ perspectives in many research fields. In drug discovery (DD), a plethora of techniques that are based on ML are being published every month. These are applied to many of the stages of the DD pipeline, from binding site characterization and prediction of ligand affinities to calculations of ADMET properties [1,2] and development and analysis of clinical trials [3,4]. Still, the ML application that has produced the largest impact on DD is possibly the application to protein structure prediction, such as RosettaFold [5] and AlphaFold2 (AF2) [6]. Using a combination of physics-based and knowledge-based energy functions, along with evolutionary information, AF2 has shown unprecedented results when predicting structures from sequence alone, leading to a dramatic increase in accuracy [6–8]. Moreover, AF2 has shown great potential in solving additional structural problems, including characterization of protein-protein interactions [9,10], protein-peptide complexes [11], and the modeling of conformational transitions for drug receptors [12].

The advances in ML paralleled by advances in more “traditional” molecular mechanics based techniques. Improved force fields [13], extended atomistic simulations using high performance computing (HPC) [14], advanced methods such as free energy perturbation (FEP) based on molecular dynamics (MD) [15] or Monte Carlo (MC) [16,17] approaches, have led to an increase in molecular modeling applications in DD by academic and industrial researchers. And these developments might even be further revalorized by their application in concert with AF2. Having access to accurate structures, as well as multiple configurational states could result in increased application of structure based DD projects, a proof of that is AlphaFold-Database [18,19], which has been widely used in many projects [20,21]. Thus, it is important to assess the performance of AF2 structures when combined with current molecular modeling methods, an aspect that has already gained significant attention elsewhere [22].

For example, in a recent publication we showed how a state of the art implementation of FEP (FEP+ from Schrödinger), when applied to AF2 structures, could produce analogous results to those when using crystal structures [23]. In that study, in order to impose more realistic prospective conditions on our benchmark experiment, we developed a custom AF2 version, named AF2_30_, where we eliminated all structural templates with >30% identity to the target protein from the training set. We observed that in most cases, ΔΔG values from AF2_30_ structures were comparable in accuracy to the corresponding calculations previously carried out using X-ray structures. Similar results, when looking at enrichment factors (EF) in docking calculations, have been recently provided by Schrödinger when using their latest induced fit approach (IFD-MD) [22]. Importantly, EF are reduced considerably, from those obtained with IFD-MD, when using simple rigid docking (Glide software and SP score) on top of AF2 structures.

In this letter we provide an analysis similar to that provided by Schrödinger [22] around the use of AF2 models for docking calculations. However, we attempt to simulate a more real-world scenario around carrying out virtual screening for a novel protein (no related protein structures available), where high-identity templates (those with identity > 30%) are removed (AF2_30_) prior to model building. In addition, we perform our calculations using out-of-the-box settings of the docking software in order to offer the point of view of an outsider user. Our results indicate that AF2 models are not suited to run rigid docking, and would require some additional treatment in order to open the cavity.

## Methods

### Dataset

A total of 10 targets from DUD-e dataset [24] were selected, including two GPCRs, two kinases, two proteases, one ion channel, one nuclear receptor, one esterase, and one lipid transport protein. The chosen systems and reference PDBs for the dataset are shown in table Table 1.

**Table 1.** Systems selected from DUD-e database. The number of active and decoy molecules correspond to the number obtained generating all possible stereoisomers with LigPrep.

| System | Uniprot ID | Reference PDB | Class | Name | Actives | Decoys |
| --- | --- | --- | --- | --- | --- | --- |
| aa2ar | P29274 | 3EML | GPCR | Adenosine A2a receptor | 2977 | 86333 |
| aces | P04058 | 1E66 | Other Enzymes | Acetylcholinesterase | 153 | 3953 |
| adrb2 | P07550 | 3NY8 | GPCR | Beta-2 adrenergic receptor | 1952 | 33925 |
| dpp4 | P27487 | 2I78 | Protease | Dipeptidyl peptidase IV | 7660 | 99580 |
| esr1 | P03372 | 1SJ0 | Nuclear Receptor | Estrogen receptor alpha | 1954 | 26215 |
| fa7 | P08709 | 1W7X | Protease | Coagulation factor VII | 305 | 14999 |
| fabp4 | P15090 | 2NNQ | Miscellaneous | Fatty acid binding protein adipocyte | 94 | 2654 |
| fak1 | Q05397 | 3BZ3 | Kinase | Focal adhesion kinase 1 | 151 | 4168 |
| gria2 | P42262 | 3KGC | Ion Channel | Glutamate receptor ionotropic, AMPA 2 | 283 | 7726 |
| mapk2 | P49137 | 3M2W | Kinase | MAP kinase-activated protein kinase 2 | 734 | 12404 |

### AF2 models

We built two different sets of AF2 structural models, AF2_30_ and AF2_100_; in both of them we used the full Multiple Sequence Analysis (MSA) obtained by default, using the full protein sequence deposited in uniprot. AF2_100_ was built with the default AF2 version, with all available templates released before 2021-12-10. AF2_30_ employs our custom AF2 version where structural templates are restricted to a 30% maximum sequence identity (see below).

For esr1, we built an antagonist-like structural model following the protocol described in Heo and Feig [12], replacing the PDB70 database by a GPCRs database in open conformation (antagonist-like) and modifying residues in the MSAs to gaps for those positions that were aligned to a structural template.

### System preparation

All structures were processed by the Protein Preparation wizard [25,26], including a hydrogen bond optimization by PROPKA at pH=7, followed by a restrained minimization (only applied on hydrogens). For AlphaFold models, first we aligned the best model (ranked_0) to the reference crystal structure in the DUD-e dataset (Table 1), in order to include the ligand.

For the ligands, in order to ensure consistent ratio of actives to decoys, we encoded the compounds into fingerprints, clustered the ligands by similarity using the Tanimoto coefficient. and extracted as many decoys centroids as necessary to establish a ratio of 1:30. All ligands were processed with LigPrep [27] to generate all possible protonation states at pH = 7.0 +/− 0.5, using Epik. Since ligands in DUD-e do not contain stereochemistry information, we verified with RDKit [28] that all possible stereoisomers were generated.

### Ligand Docking

To perform ligand docking we used Glide [29] from the Schrödinger suite [30]. Using the ligand from the reference PDB, we selected the center of the docking grid box to the central atom (the one closest to the ligands center-of-mass (COM)). Then we modified the inner box (following best practices at Nostrum Biodiscovery and by using advanced settings) to cover the volume of the entire ligand while the dimensions of the outer box were generated automatically. All SP docking parameters were set to default. Only the best scoring glide pose was considered in the following analysis of docking results.

The EF was computed with the enrichment factor calculator provided by Schrödinger, enrichment.py. FPocket [31] was used to compute the active site volumes, where we used the -r option to specify the ligand and have a better approximation of the binding site (BS) volume.

### AF2 customization

Our AF2 customization has been described in detail in our recent FEP study [23]. Briefly we systematically removed all template structures above 30% sequence identity from the database used to build the models. Our version is now capable of removing either structural templates, or sequences, or both, from the AF2 database, based on a user-defined sequence identity threshold.

## Results and Discussion

Table 2 shows the ligand root mean square distance (RMSD) values for the docking of the reference PDB ligand into its cognate crystal structure. Clearly all but one system give very low RMSD values in the X-ray self docking experiment. Only the adenosine A2a receptor introduces a high RMSD value, due to the absence of two crystal waters in the target model (as per default we remove all water molecules that could hinder ligand binding). If those water molecules are included in the docking grid, the native pose is retrieved; in addition a soft docking exercise retrieves a native-like pose, although not as the highest ranked solution. As expected, much higher RMSD values are found when using the AF2 models (both at 100% and 30% sequence cutoff), which reflect the difficulties in performing docking into apo-like structures. In most cases AF2 models introduce changes that partially fill the (empty) pocket space. Most of these changes occur at the side chain level, including the movement of Asp113 and Asn312 in adrb2 (Figure 1), for example. However larger changes are also observed such as the rearrangement of the 423-426 loop in fa7 or the agonist-like AF2 pose for esr1. In general we observe similar behavior for AF2_100_ and AF2_30_, reflecting that AF2 predicts good structures even in absence of closely related structural templates. We observe, however, that for those two systems where AF2_100_ provides an RMSD <2 Å, adrb2 and mapk2, AF2_30_ provides worst results; the most drastic case being adrb2 where AF2_30_ fails in providing near crystal poses. This worsening of the results is more evident when performing a more general screening exercise (see below).

**Figure 1.**
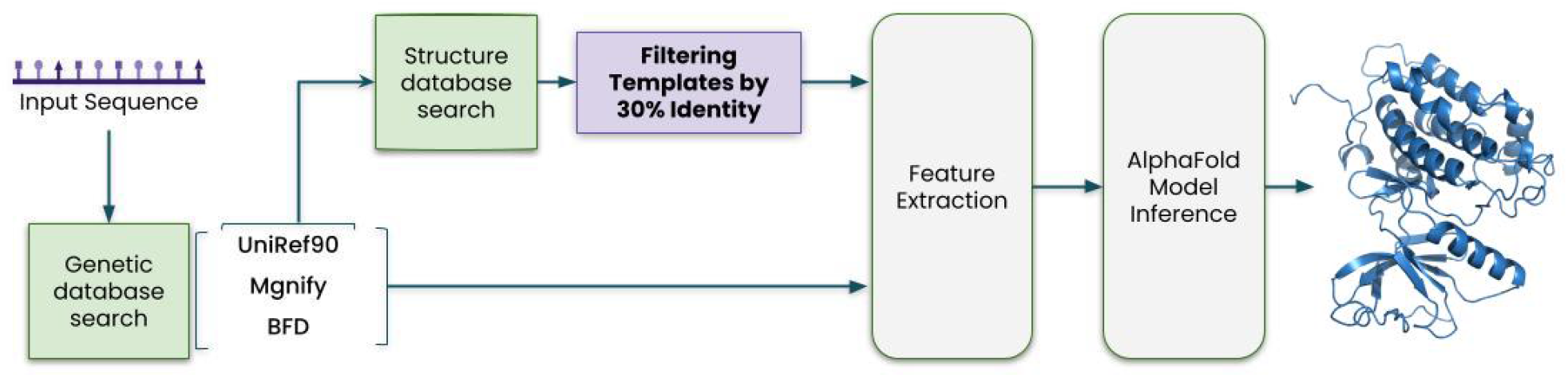
AF2_30_ workflow used in this work, where we removed all PDB templates with sequence above 30% to the target, leaving all MSA information intact.

**Table 2.**
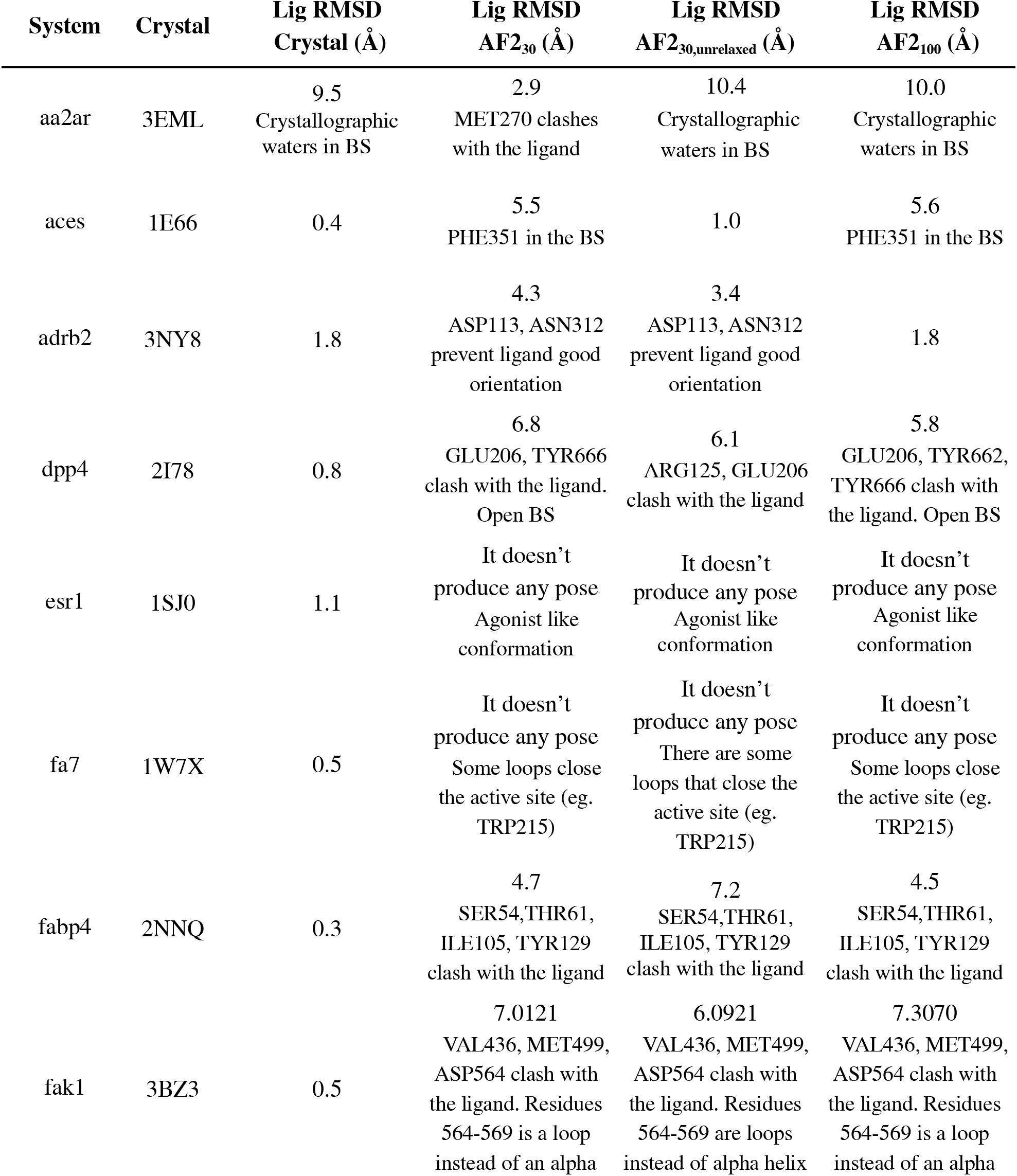
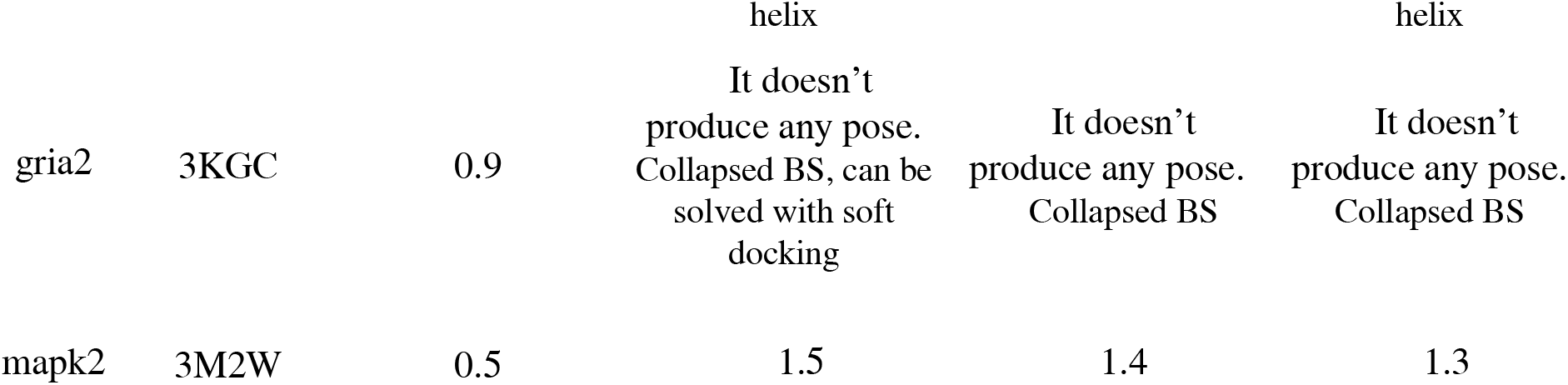
Self docking of DUDE’s reference PDB, showing all atom RMSD of the cognate ligand after Glide docking. For those cases where there was no pose returned by Glide or where the RMSD was above 2 Å, we provide a short additional explanation.

An example of a subtle but significant active site collapse is shown in Figure 2. Several side chains in adrb2 slightly close the space, enough to force the ligand to adopt a different conformation when docked. The active site volume as computed with Fpocket reduces from 901 Å^3^ in the crystal to 723 Å^3^ in AF2_30_ (see all data set values in Supporting Information Table S1). Interestingly, such a collapse is partly reduced if omitting the last AF2 minimization, which is performed as a final relaxation phase using the AMBER force field in order to alleviate steric clashes (notice that AF2 provides an unrelaxed output structure). Indeed when performing self docking with unrelaxed AF2_30_ we obtain better RMSD in most of the values. However, for two of the systems, aa2ar and fabp4, for which the results are already bad, unrelaxed predictions get worse. In these systems we only observe tiny structural variations upon relaxation which translate into small relative changes in the volume (Table S1).

**Figure 2.**
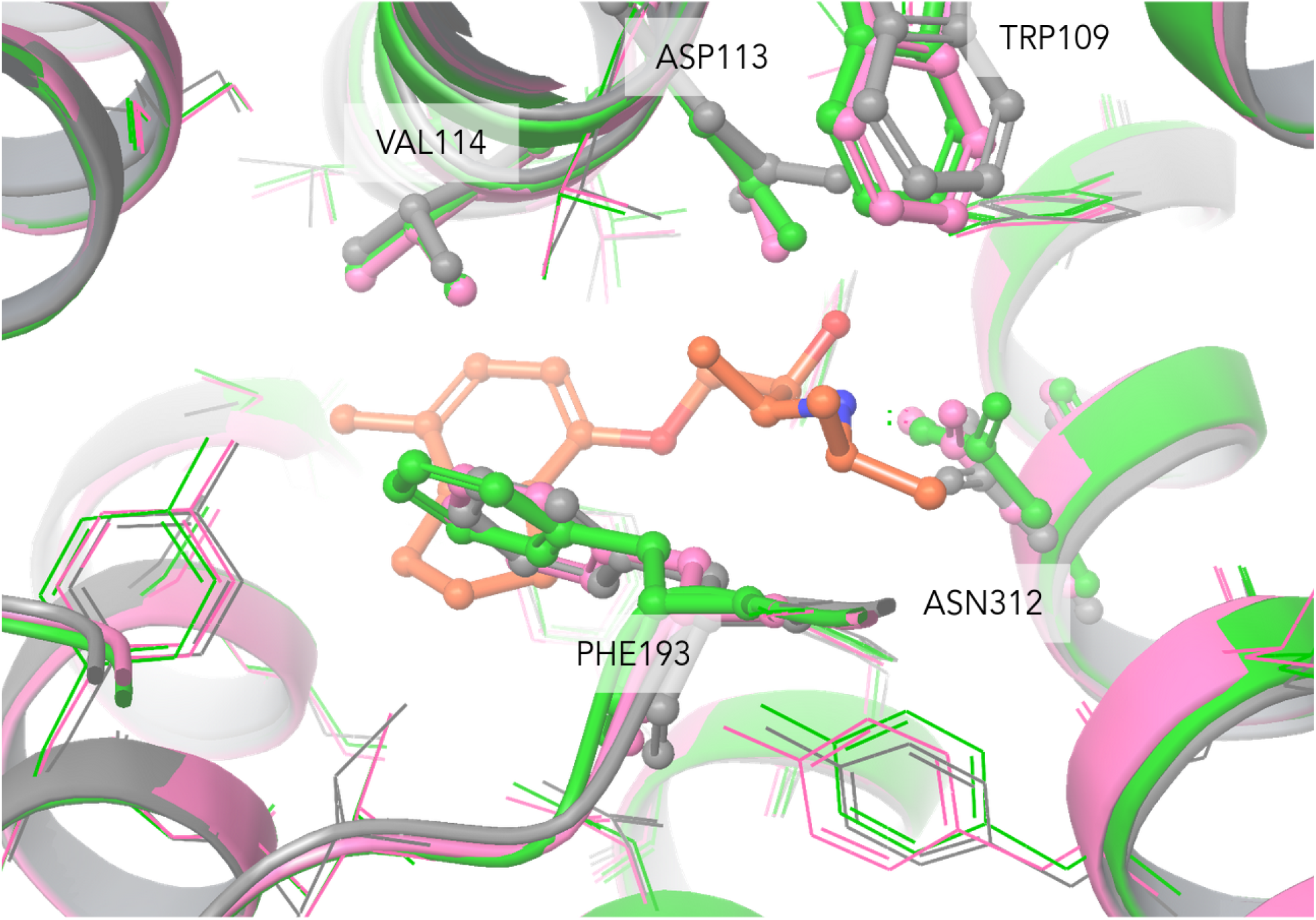
Adrb2 X-ray (gray), AF2_30_ (green) and AF2_100_ (pink) binding site structures after alpha carbon protein superposition. The X-ray ligand is also shown in element-colored ball and stick.

AF2 difficulties in accurately reproducing holo features in the binding pocket are also translated into significantly poorer EF; table 3 shows EF 1% values for all AF2 models. Please refer to supporting information for additional ROC and EF values, which follow the same trend. Moreover, if performing a more realistic analysis, as the one provided with AF2_30_ models, the results are quite disappointing. In fabp4, fak1 and gria2, for example, we observe drastic reductions of EF. In gria2 there is a large reduction of the active site volume (Table S1 and Figure S1.I); notice how the AF_100_ model has a ~30% reduction when comparing it to the AF_30_ one, resulting in a significantly worse EF. In fabp4 and fak1, while volumes are similar, AF2 spatial disposition in the active sites have large perturbations when comparing it to the crystal: side chain clashes in fabp4 and an alpha helix predicted as a loop in fak1 (Figure S1). Interestingly, in one system, aa2ar, we observe increased EFs for AF2 models. When inspecting the predicted binding site volumes (Table S1), this is the only system where we observe a significant increase for the modeled targets.

**Table 3.** 1% Enrichment factors for all the dataset.

| System | Crystal | AF2<sub>30</sub> | AF2<sub>30,unrelaxed</sub> | AF2<sub>100</sub> |
| --- | --- | --- | --- | --- |
| aa2ar | 8.5 | 14.0 | 12.0 | 10.0 |
| aces | 0.7 | 0.0 | 0.0 | 0.7 |
| adrb2 | 9.0 | 5.9 | 4.2 | 6.6 |
| dpp4 | 6.1 | 2.2 | 5.2 | 1.9 |
| esr1 | 14.0 | 6.2 | 6.2 | 10.0 |
| esr1 (open-conf) | 14.0 | - | - | 13.0 |
| fa7 | 13.0 | 9.0 | 11.0 | 13.0 |
| fabp4 | 30.0 | 5.6 | 8.7 | 9.7 |
| fak1 | 26.0 | 3.3 | 3.3 | 5.3 |
| gria2 | 20.0 | 13.0 | 11.0 | 6.4 |
| mapk2 | 15.0 | 14.0 | 7.1 | 13.0 |

The reduced performance of AF2 models when compared to X-ray holo structures is aligned with the recent study of Schrödinger [19], and leads us to conclude that performing a rigid receptor docking exercise using out-of-the-box AF2 models is not an ideal scenario. A post processing step, including some pocket opening/readjustment seems necessary to reach practically useful EF values. For example, the IFD-MD technology largely improved EF factors in the Schrödinger study. A cheaper (faster) alternative could be to use AF2 models before the final minimization. Since the worse AF2 models performance partly originates from the binding site volume reduction (collapse), and because unrelaxed AF2 models, which are provided as well by the default application, slightly increase this volume, we explore this alternative. While results on self docking were promising, EFs do not corroborate a valid trend.

## Conclusion

Overall, our study indicates that the use of Alphafold2 models for virtual screening has to be performed with a fair amount of caution. Models tend to be representative of apo structures without explicit water molecules, introducing a significant collapse of the binding sites when compared with holo crystals. Thus, their use in molecular modeling activities requires additional care. For the initial step, we recommend comparing the top four templates used by AF2, superposing and analyzing potential differences in the binding site. For the next step, before a large scale virtual screening campaign, it would be ideal to run some induced fit application (such as PELE [32] or IFD-MD [22]) using known inhibitors, aiming for a partial opening/accomodation of the active site residues. If the binding site still appears to be collapsed, we recommend then running extensive MD simulations to potentially open these pockets in order to accommodate ligands. Still, observing these recommendations and with a little knowledge of the target, AF2 could be a good tool for target generation applied to virtual screening campaigns in absence of crystal structures.

## Author Contributions

The manuscript was written through contributions of all authors. All authors have given approval to the final version of the manuscript.

## ABBREVIATIONS

ML: Machine Learning;
DD: Drug Discovery;
AF2: AlphaFold2;
HPC: High Performance Computing;
FEP: Free Energy Perturbation;
MD: Molecular Dynamics;
MC: Monte Carlo;
EF: Enrichment Factor;
IDF-MD: Schrödinger Induced fit approach;
aa2ar: Adenosine receptor A2a;
aces: Acetylcholinesterase;
adrb2: Beta-2 adrenergic receptor;
dpp4: Dipeptidyl peptidase IV;
esr1: Estrogen receptor alpha;
fa7: Coagulation factor VII;
fabp4: Fatty acid binding protein adipocyte;
fak1: Focal adhesion kinase 1;
gria2: Glutamate receptor ionotropic AMPA 2;
mapk2: MAP kinase-activated protein kinase 2;
MSA: Multiple Sequence Alignment;
COM: Center-Of-Mass;
BS: Binding Site;
RMSD: Root Mean Square Distance.

## DATA AVAILABILITY

All AF2_30_ and AF2_100_ models used as input for Glide docking, along with ligand poses have been uploaded as supplementary information

## CODE AVAILABILITY

Our custom AlphaFold2 implementation to remove sequences and/or templates is published on GitHub under Apache License 2.0: https://github.com/hemahecodes/alphafold.

