## Supplementary Information for "Are Deep Learning Structural Models Sufficiently Accurate for Virtual Screening? Application of Docking Algorithms to AlphaFold2 Predicted Structures"

**Table S1**. BS Volumes computed with FPocket and visual inspection of the models.

| System | Crystal | $AF2_{30}$ | $AF2_{30}$ unrelaxed | $AF2_{100}$ | Visual inspection |
|---|---|---|---|---|---|
| aa2ar | 1559.8 | 1829.4 | 1952.1 | 1728.7 | TRP246, MET270, HIS264 invading BS in AF models. Some alpha helices (eg. 167-171, 268-281) in the crystal are slightly moved towards the BS. |
| aces | 603.6 | 545.8 | 526.7 | 511.5 | PHE351 invading BS, prevents ligand is well oriented |
| adrb2 | 901.5 | 723.8 | 824.4 | 757.1 | ASP113, ASN312 prevent that the ligand is well oriented. TRP109, VAL114, PHE193 are occupying the BS |
| dpp4 | 1693.3 | 1779.1 | 1943.2 | 1733.5 | Secondary structure of the crystal seems to be slightly collapsed to the BS respect to AF models (eg. Alpha Helix 630-642, loop 654-656) |
| esr1 | 1431.3 | 581.9 | 616.9 | 555.8 | Closed conformation (agonist like conformation) |
| esr1 open-conf | 1431.3 | - | - | 833.0 | Open conformation (antagonist like conformation) |
| fa7 | 2526.3 | 1962.3 | 1939.4 | 2168.8 | There are loops that close the active site (resnums 400-403, 423-426) |
| fabp4 | 965.8 | 961.4 | 1034.2 | 880.01 | SER54,THR61, ILE105, TYR129 badly oriented (clash with the ligand) |

| | | | | | |
|---|---|---|---|---|---|
| fak1 | 1977.3 | 2022.7 | 1873.1 | 1896.9 | VAL436, MET499, ASP564 clash with the ligand. 564-569 are loops instead of alpha helix |
| gria2 | 1676.1 | 759.0 | 852.1 | 516.4 | Alpha helices in the BS (resnum 673-675, 706-707). Loops in the BS (resnum 421-473, 498-503) |
| mapk2 | 2095.6 | 1548.3 | 1539.6 | 1548.3 | ASP142 pointing to BS. Slightly more collapsed BS (LEU70, GLU139, CYS140, LEU141), Moved Alpha Helix (resnum 69-73) |

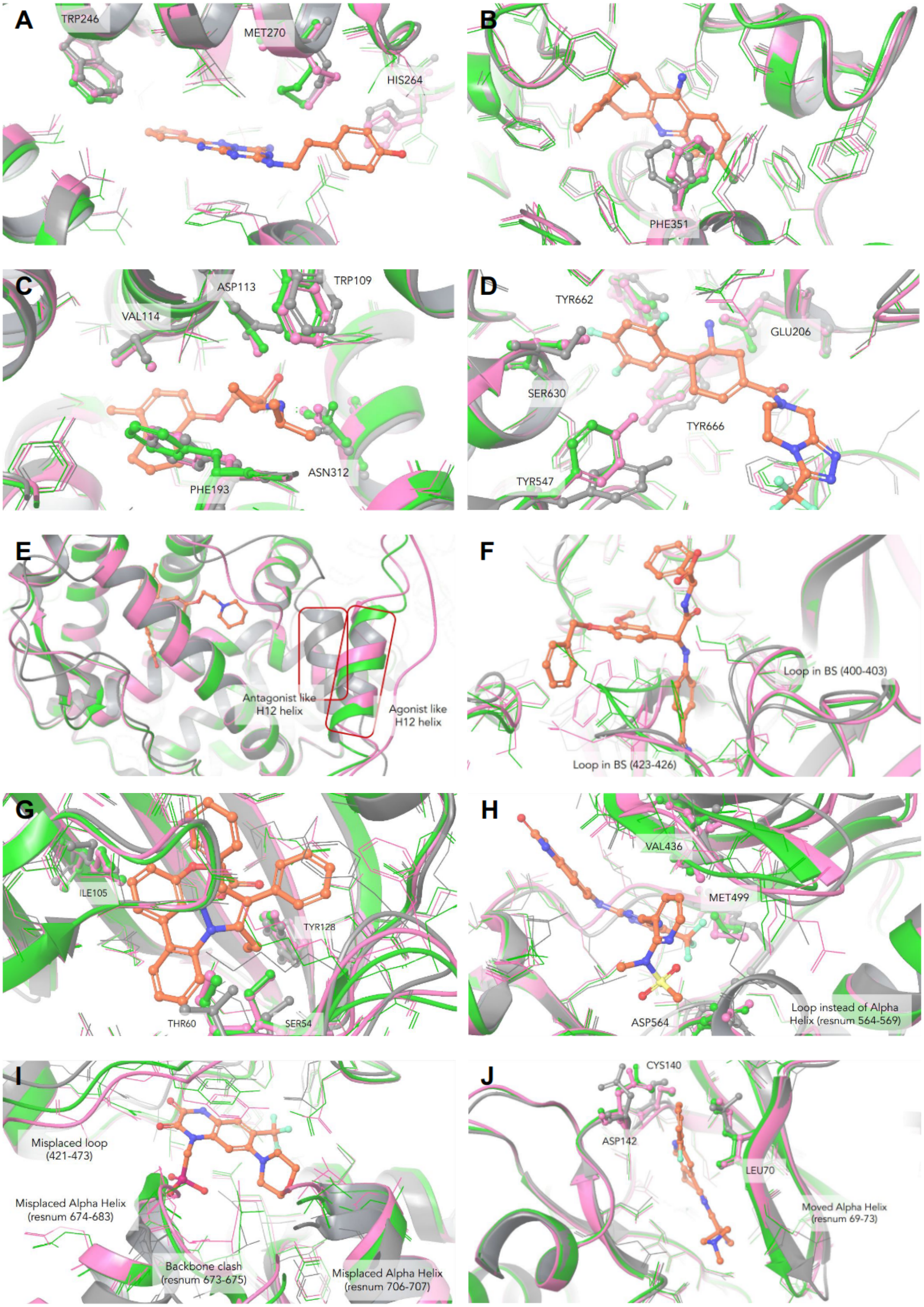


**Figure S1**. BS 3D structure of the dataset after alpha carbon protein superposition. A) aa2ar, B)

aces, C) adrb2, D) dpp4, E) esr1, F) fa7, G) fabp4, H) fak1, I) gria2, J) mapk2. The X-ray reference crystal for each system is in gray, $AF2_{30}$ models are in green while $AF2_{100}$ models are in pink. The X-ray ligand is also shown in element-colored ball and stick.

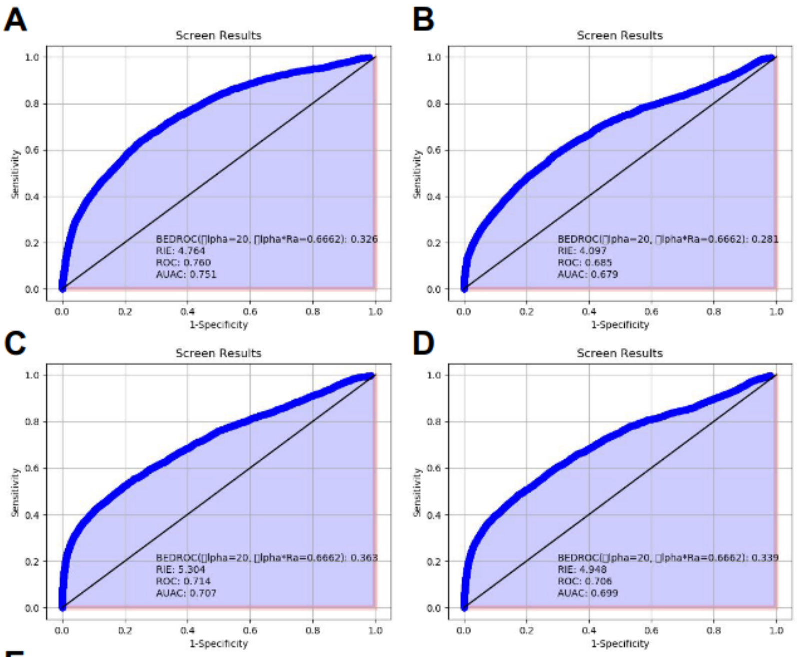


**E**

| %EF | Top 1% | Top 2% | Top 5% | Top 10% | Top 20% |
|---|---|---|---|---|---|
| Crystal | 8.5 | 7.8 | 5.8 | 4.0 | 2.8 |
| $AF2_{100}$ | 10.0 | 7.5 | 4.6 | 3.2 | 2.3 |
| $AF2_{30}$ | 14.0 | 11.0 | 6.3 | 4.0 | 2.6 |
| $AF2_{30, unrelaxed}$ | 12.0 | 9.7 | 5.8 | 3.8 | 2.5 |

**F**

| %Actives in... | Top 1% | Top 2% | Top 5% | Top 10% | Top 20% |
|---|---|---|---|---|---|
| Crystal | 8.5 | 15.6 | 29.1 | 40.2 | 55.1 |
| $AF2_{100}$ | 10.2 | 15.1 | 23.1 | 32.4 | 46.6 |
| $AF2_{30}$ | 14.2 | 21.6 | 31.3 | 39.8 | 51.1 |
| $AF2_{30. unrelaxed}$ | 12.2 | 19.3 | 29.2 | 38.2 | 49.7 |

**Figure S2**. Analysis of glide results for aa2ar system. ROC curves of A) the reference crystal, B) $AF2_{100}$, C) $AF2_{30}$ and D) $AF2_{30,unrelaxed}$. E) EF when selecting the top N% ranked ligands. F) Percentage of actives in top N% of results.

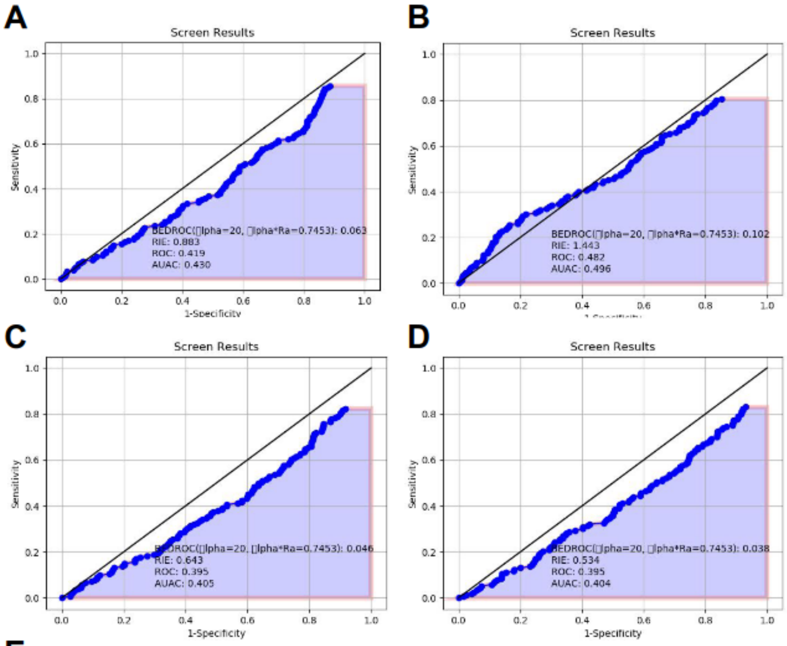


**E**

| %EF | Top 1% | Top 2% | Top 5% | Top 10% | Top 20% |
|---|---|---|---|---|---|
| Crystal | 0.7 | 1.3 | 0. | 0.8 | 0.8 |
| $AF2_{100}$ | 0.7 | 1.6 | 1.4 | 1.4 | 1.3 |
| $AF2_{30}$ | 0.0 | 0.0 | 0.7 | 0.7 | 0.7 |
| $AF2_{30,\ unrelaxed}$ | 0.0 | 0.3 | 0.5 | 0.5 | 0.7 |

**F**

| %Actives in... | Top 1% | Top 2% | Top 5% | Top 10% | Top 20% |
|---|---|---|---|---|---|
| Crystal | 0.7 | 2.6 | 4.6 | 7.8 | 15.7 |
| $AF2_{100}$ | 0.7 | 3.3 | 7.2 | 13.7 | 26.8 |
| $AF2_{30}$ | 0.0 | 0.0 | 3.3 | 7.2 | 14.4 |
| $AF2_{30,\ unrelaxed}$ | 0 | 0.7 | 2.6 | 5.2 | 13.1 |

**Figure S3**. Analysis of glide results for aces system. ROC curves of A) the reference crystal, B) $AF2_{100}$, C) $AF2_{30}$ and D) $AF2_{30,unrelaxed}$. E) EF when selecting the top N% ranked ligands. F) Percentage of actives in top N% of results.

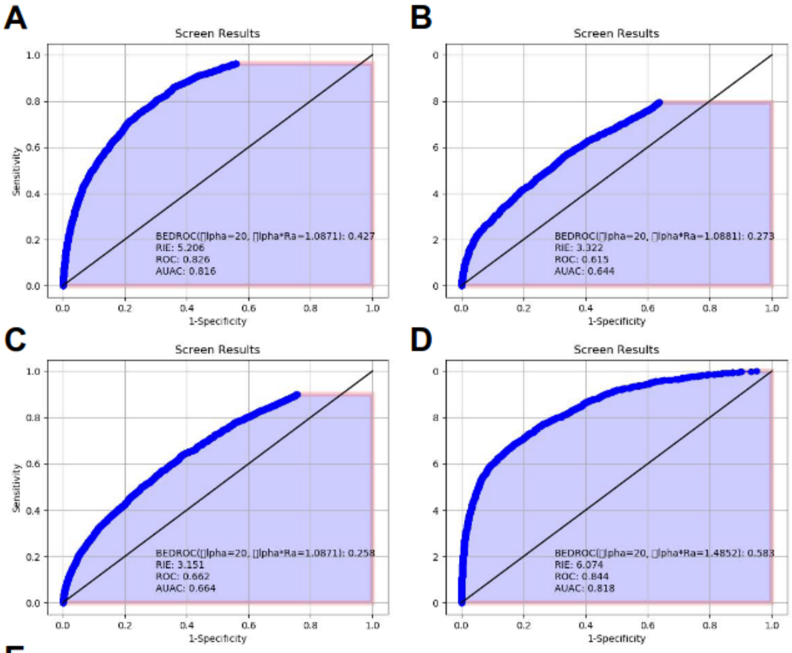


**E**

| %EF | Top 1% | Top 2% | Top 5% | Top 10% | Top 20% |
|---|---|---|---|---|---|
| Crystal | 9.0 | 8.1 | 6.1 | 4.6 | 3.2 |
| $AF2_{100}$ | 6.6 | 5.7 | 4.0 | 2.7 | 2.0 |
| $AF2_{30}$ | 5.9 | 4.8 | 3.6 | 2.8 | 2.1 |
| $AF2_{30,\ unrelaxed}$ | 4.2 | 4.1 | 3.3 | 2.6 | 2.1 |

**F**

| %Actives in... | Top 1% | Top 2% | Top 5% | Top 10% | Top 20% |
|---|---|---|---|---|---|
| Crystal | 9.0 | 16.2 | 30.2 | 46.1 | 64.3 |
| $AF2_{100}$ | 6.6 | 11.3 | 20.0 | 27.3 | 40.7 |
| $AF2_{30}$ | 5.9 | 9.7 | 17.8 | 27.5 | 41.0 |
| $AF2_{30,\ unrelaxed}$ | 4.2 | 8.2 | 16.4 | 25.8 | 41.6 |

**Figure S4**. Analysis of glide results for adrb2 system. ROC curves of A) the reference crystal, B) $AF2_{100}$, C) $AF2_{30}$ and D) $AF2_{30,unrelaxed}$. E) EF when selecting the top N% ranked ligands. F) Percentage of actives in top N% of results.

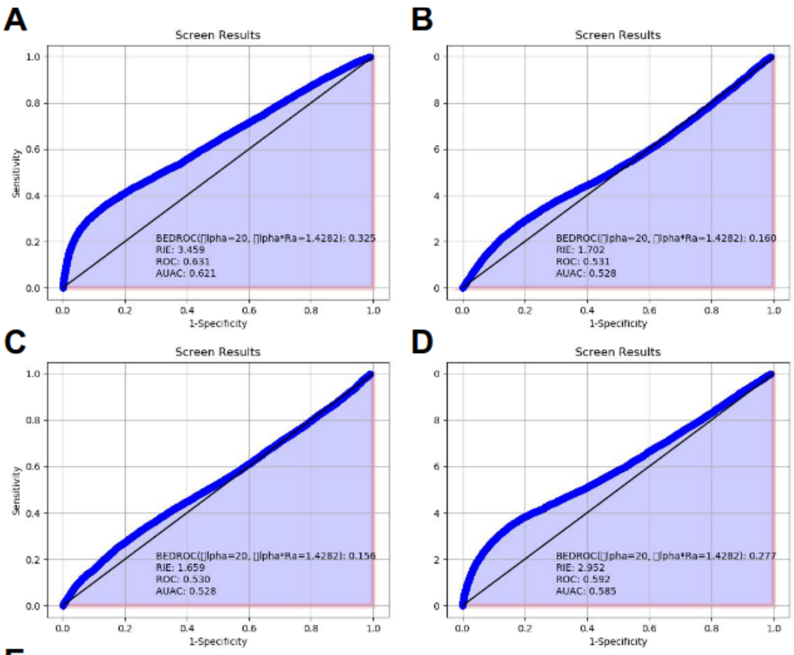


**E**

| %EF | Top 1% | Top 2% | Top 5% | Top 10% | Top 20% |
|---|---|---|---|---|---|
| Crystal | 6.1 | 5.5 | 4.2 | 3.0 | 2.0 |
| $AF2_{100}$ | 1.9 | 1.9 | 1.8 | 1.7 | 1.4 |
| $AF2_{30}$ | 2.2 | 2.0 | 1.9 | 1.5 | 1.3 |
| $AF2_{30,\ unrelaxed}$ | 5.2 | 4.5 | 3.5 | 2.6 | 1.9 |

**F**

| %Actives in... | Top 1% | Top 2% | Top 5% | Top 10% | Top 20% |
|---|---|---|---|---|---|
| Crystal | 6.1 | 11.0 | 21.1 | 29.9 | 39.9 |
| $AF2_{100}$ | 1.9 | 3.7 | 9.2 | 17.0 | 28.4 |
| $AF2_{30}$ | 2.2 | 4.0 | 9.3 | 15.2 | 26.8 |
| $AF2_{30,\ unrelaxed}$ | 5.2 | 9.0 | 17.5 | 26.4 | 37.3 |

**Figure S5**. Analysis of glide results for dpp4 system. ROC curves of A) the reference crystal, B) $AF2_{100}$, C) $AF2_{30}$ and D) $AF2_{30,unrelaxed}$. E) EF when selecting the top N% ranked ligands. F) Percentage of actives in top N% of results.

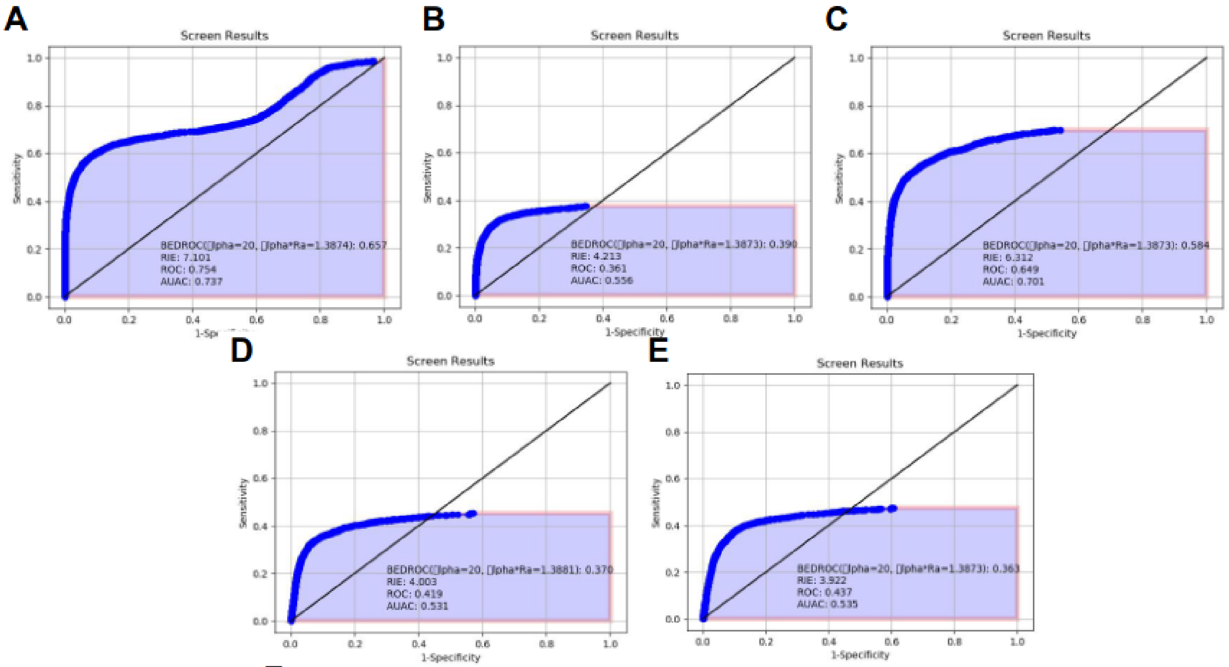


**F**

| %EF | Top 1% | Top 2% | Top 5% | Top 10% | Top 20% |
|---|---|---|---|---|---|
| Crystal | 14.0 | 13.0 | 9.1 | 5.7 | 3.2 |
| $AF2_{100}$ | 10.0 | 8.5 | 5.2 | 3.2 | 1.8 |
| $AF2_{OpenConf}$ | 13.0 | 11.0 | 8.1 | 5.1 | 3.0 |
| $AF2_{30}$ | 6.2 | 6.4 | 5.3 | 3.4 | 2.0 |
| $AF2_{30,\ unrelaxed}$ | 6.2 | 5.9 | 5.1 | 3.5 | 2.1 |

**G**

| %Actives in... | Top 1% | Top 2% | Top 5% | Top 10% | Top 20% |
|---|---|---|---|---|---|
| Crystal | 14.3 | 26.4 | 45.7 | 56.8 | 64.0 |
| $AF2_{100}$ | 6.1 | 11.8 | 24.4 | 35.2 | 41.8 |
| $AF2_{OpenConf}$ | 13.0 | 22.2 | 40.6 | 51.1 | 59.2 |
| $AF2_{30}$ | 6.1 | 12.8 | 26.5 | 34.3 | 39.7 |
| $AF2_{30,\ unrelaxed}$ | 6.1 | 11.8 | 24.4 | 35.2 | 41.8 |

**Figure S6**. Analysis of glide results for esr1 system. ROC curves of A) the reference crystal, B) $AF2_{100}$, C) $AF2_{100,OpenConf}$, D) $AF2_{30}$ and E) $AF2_{30,unrelaxed}$. F) EF when selecting the top N% ranked ligands. G) Percentage of actives in top N% of results.

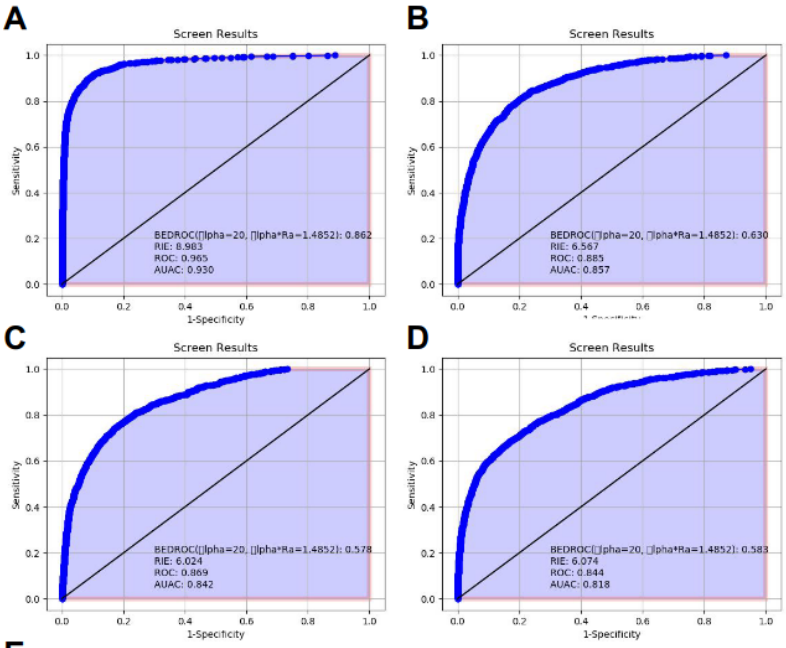


**E**

| %EF | Top 1% | Top 2% | Top 5% | Top 10% | Top 20% |
|---|---|---|---|---|---|
| Crystal | 13.0 | 13.0 | 12.0 | 8.3 | 4.7 |
| $AF2_{100}$ | 12.9 | 21.9 | 39.2 | 57.0 | 74.9 |
| $AF2_{30}$ | 9.0 | 8.6 | 7.5 | 5.4 | 3.6 |
| $AF2_{30,\ unrelaxed}$ | 11.0 | 9.9 | 7.4 | 5.4 | 3.4 |

**F**

| %Actives in... | Top 1% | Top 2% | Top 5% | Top 10% | Top 20% |
|---|---|---|---|---|---|
| Crystal | 13.2 | 25.5 | 58.1 | 82.5 | 93.5 |
| $AF2_{100}$ | 12.9 | 21.9 | 39.2 | 57.0 | 74.9 |
| $AF2_{30}$ | 9.0 | 17.3 | 37.7 | 54.2 | 72.2 |
| $AF2_{30,\ unrelaxed}$ | 10.9 | 19.8 | 37.0 | 53.6 | 67.5 |

**Figure S7**. Analysis of glide results for fa7 system. ROC curves of A) the reference crystal, B) $AF2_{100}$, C) $AF2_{100,OpenConf}$, D) $AF2_{30}$ and E) $AF2_{30,unrelaxed}$. F) EF when selecting the top N% ranked ligands. G) Percentage of actives in top N% of results.

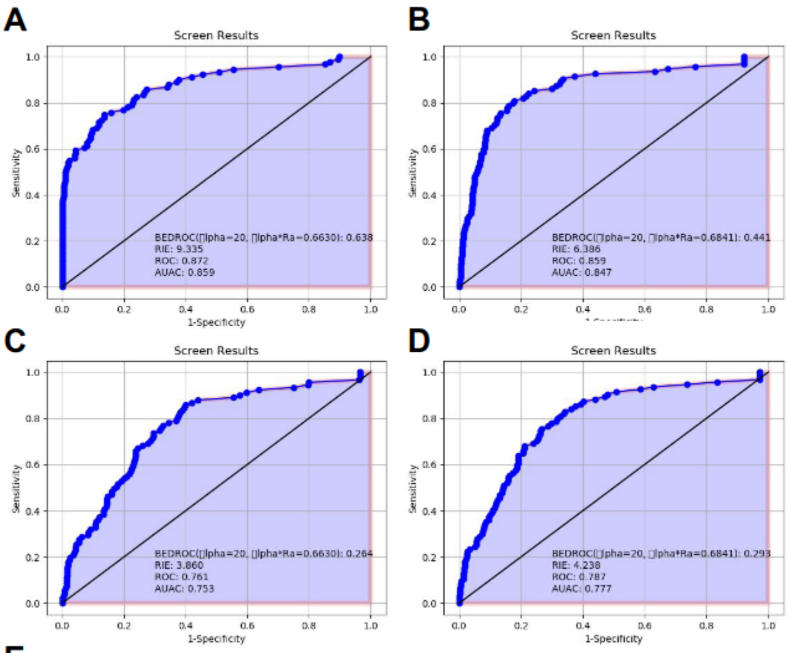


**E**

| %EF | Top 1% | Top 2% | Top 5% | Top 10% | Top 20% |
|---|---|---|---|---|---|
| Crystal | 30.0 | 22.0 | 11.0 | 6.3 | 3.8 |
| AF2$_{100}$ | 9.7 | 10.0 | 6.6 | 6.0 | 4.0 |
| AF2$_{30}$ | 5.6 | 6.0 | 4.8 | 3.2 | 2.6 |
| AF2$_{30,\ unrelaxed}$ | 8.7 | 6.4 | 4.7 | 3.5 | 2.9 |

**F**

| %Actives in... | Top 1% | Top 2% | Top 5% | Top 10% | Top 20% |
|---|---|---|---|---|---|
| Crystal | 29.7 | 42.9 | 54.9 | 62.6 | 75.8 |
| AF2$_{100}$ | 9.6 | 20.2 | 33.0 | 59.6 | 80.9 |
| AF2$_{30}$ | 5.5 | 11.0 | 24.2 | 31.9 | 51.6 |
| AF2$_{30,\ unrelaxed}$ | 8.5 | 12.8 | 23.4 | 35.1 | 57.4 |

**Figure S8**. Analysis of glide results for fabp4 system. ROC curves of A) the reference crystal, B) AF2$_{100}$, C) AF2$_{100,OpenConf}$, D) AF2$_{30}$ and E) AF2$_{30,unrelaxed}$. F) EF when selecting the top N% ranked ligands. G) Percentage of actives in top N% of results.

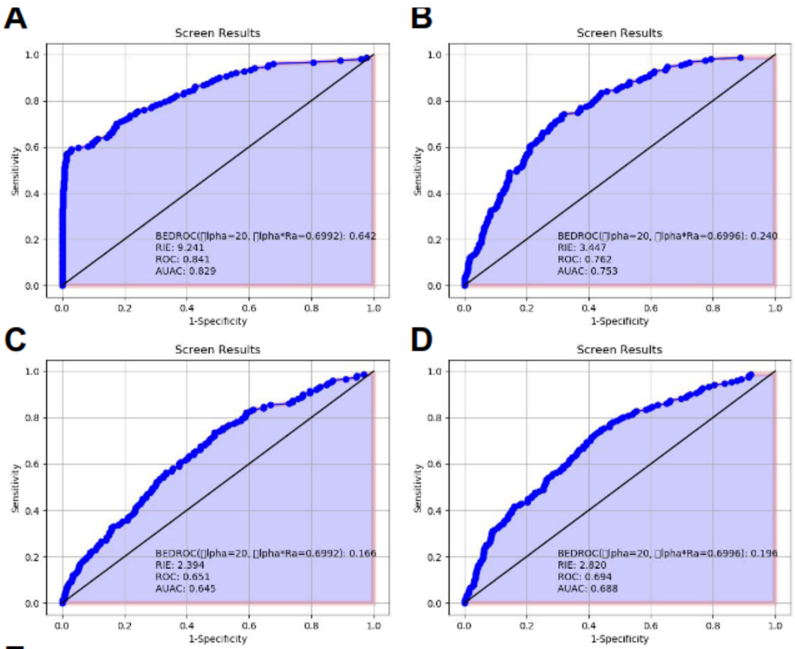


E

| %EF | Top 1% | Top 2% | Top 5% | Top 10% | Top 20% |
|---|---|---|---|---|---|
| Crystal | 26.0 | 21.0 | 12.0 | 6.0 | 3.5 |
| $AF2_{100}$ | 5.3 | 6.0 | 3.3 | 3.2 | 2.6 |
| $AF2_{30}$ | 3.3 | 3.7 | 2.6 | 2.2 | 1.7 |
| $AF2_{30,\ unrelaxed}$ | 3.3 | 3.3 | 3.3 | 3.1 | 2.2 |

F

| %Actives in... | Top 1% | Top 2% | Top 5% | Top 10% | Top 20% |
|---|---|---|---|---|---|
| Crystal | 25.8 | 42.4 | 58.9 | 59.6 | 70.2 |
| $AF2_{100}$ | 5.3 | 11.9 | 16.6 | 32.5 | 52.3 |
| $AF2_{30}$ | 3.3 | 7.3 | 13.2 | 21.9 | 34.4 |
| $AF2_{30,\ unrelaxed}$ | 3.3 | 3.7 | 2.6 | 2.2 | 1.7 |

**Figure S9**. Analysis of glide results for fak1 system. ROC curves of A) the reference crystal, B) $AF2_{100}$, C) $AF2_{100,OpenConf}$, D) $AF2_{30}$ and E) $AF2_{30,unrelaxed}$. F) EF when selecting the top N% ranked ligands. G) Percentage of actives in top N% of results.

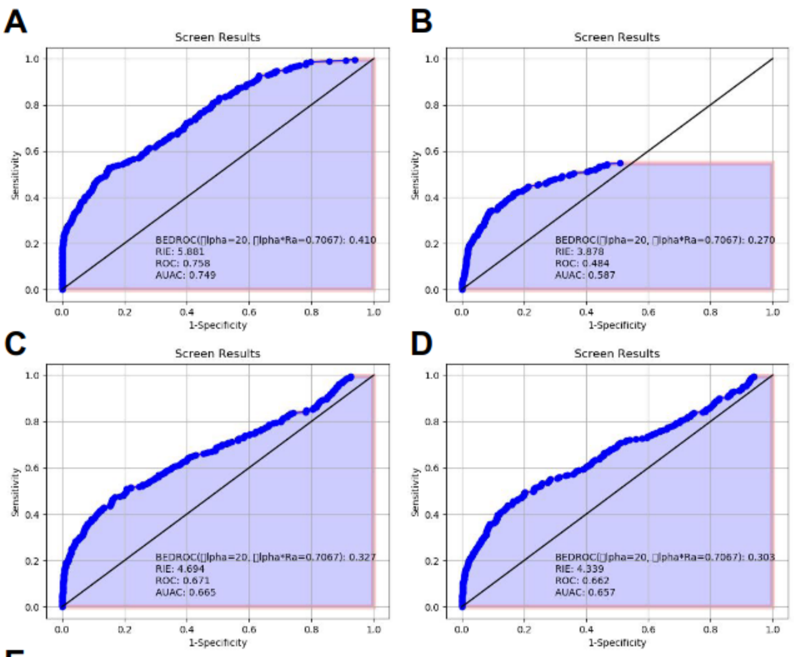


**E**

| %EF | Top 1% | Top 2% | Top 5% | Top 10% | Top 20% |
|---|---|---|---|---|---|
| Crystal | 20.0 | 13.0 | 6.7 | 4.2 | 2.7 |
| $AF2_{100}$ | 6.4 | 6.9 | 4.7 | 3.4 | 2.1 |
| $AF2_{30}$ | 13.0 | 9.6 | 5.5 | 3.7 | 2.4 |
| $AF2_{30,\ unrelaxed}$ | 11.0 | 8.3 | 4.9 | 3.6 | 2.3 |

**F**

| %Actives in... | Top 1% | Top 2% | Top 5% | Top 10% | Top 20% |
|---|---|---|---|---|---|
| Crystal | 20.1 | 25.1 | 33.2 | 42.0 | 54.1 |
| $AF2_{100}$ | 8.1 | 17.3 | 24.0 | 34.3 | 43.1 |
| $AF2_{30}$ | 13.4 | 19.1 | 27.6 | 36.4 | 47.7 |
| $AF2_{30,\ unrelaxed}$ | 11.3 | 16.6 | 24.4 | 35.7 | 46.6 |

**Figure S10**. Analysis of glide results for gria2 system. ROC curves of A) the reference crystal, B) $AF2_{100}$, C) $AF2_{100,OpenConf}$, D) $AF2_{30}$ and E) $AF2_{30,unrelaxed}$. F) EF when selecting the top N% ranked ligands. G) Percentage of actives in top N% of results.

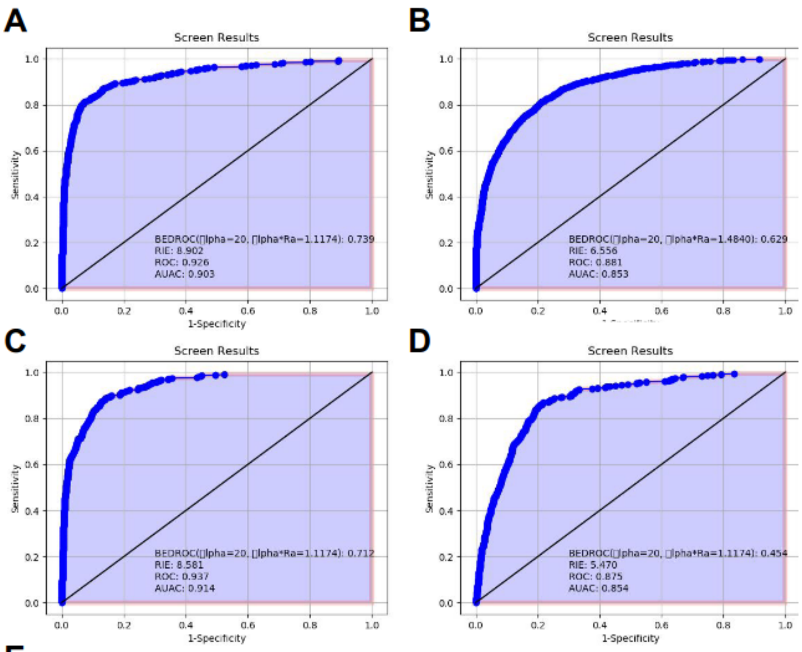


**E**

| %EF | Top 1% | Top 2% | Top 5% | Top 10% | Top 20% |
|---|---|---|---|---|---|
| Crystal | 15.0 | 14.0 | 11.0 | 7.9 | 4.4 |
| $AF2_{100}$ | 13.0 | 12.0 | 11.0 | 7.5 | 4.4 |
| $AF2_{30}$ | 14.0 | 14.0 | 11.0 | 7.2 | 4.5 |
| $AF2_{30,\ unrelaxed}$ | 7.1 | 7.4 | 6.2 | 5 | 3.9 |

**F**

| %Actives in... | Top 1% | Top 2% | Top 5% | Top 10% | Top 20% |
|---|---|---|---|---|---|
| Crystal | 15.0 | 28.1 | 56.3 | 78.7 | 88.6 |
| $AF2_{100}$ | 12.5 | 24.0 | 54.8 | 75.5 | 86.9 |
| $AF2_{30}$ | 13.6 | 27.0 | 55.6 | 71.9 | 89.6 |
| $AF2_{30,\ unrelaxed}$ | 7.1 | 14.7 | 31.1 | 50.4 | 77.1 |

**Figure S11**. Analysis of glide results for mapk2 system. ROC curves of A) the reference crystal, B) $AF2_{100}$, C) $AF2_{100,OpenConf}$, D) $AF2_{30}$ and E) $AF2_{30,unrelaxed}$. F) EF when selecting the top N% ranked ligands. G) Percentage of actives in top N% of results.
